# Seasonal blood-brain barrier plasticity links environmental cues to migratory behavior in monarch butterflies

**DOI:** 10.64898/2026.01.16.699996

**Authors:** Aldrin B. Lugena, Kayla M. Goforth, Vinaya Shetty, Ying Zhang, Antonio González-Rodríguez, M. Isabel Ramírez, Christine Merlin

## Abstract

Seasonal migration requires animals to reversibly switch behavioral states in response to environmental cues, yet the molecular and cellular mechanisms underlying these transitions remain poorly understood. Monarch butterflies provide a powerful model, as Eastern North American populations undergo a long-distance southward migration in the fall followed by a cold-dependent reversal in flight orientation after overwintering each spring. Here, we show that cold exposure induces coordinated transcriptional changes in the monarch brain, marked by attenuated integrin-mediated extracellular matrix (ECM) signaling at the blood-brain barrier (BBB). Cold-exposed monarchs exhibit increased penetration of fluorescent markers into the brain, consistent with increased BBB permeability. Notably, our findings align with previous genomic evidence identifying *collagen type IV alpha 1*, a major ECM component, as a locus under divergent selection between migratory and non-migratory populations. Together, these results implicate seasonal modulation of BBB permeability as a mechanism linking environmental temperature to plastic migratory behavior and identify the BBB as a dynamic interface through which seasonal cues may reprogram neural function and behavior.

## Introduction

Seasonal changes in animal behavior, such as migration, hibernation, and breeding, reflect finely tuned adaptations to predictable environmental fluctuations. These behaviors are orchestrated by internal mechanisms that sense and translate external cues such as photoperiod, temperature, and resource availability into appropriate physiological and behavioral responses. However, the mechanisms by which seasonal environmental signals trigger long-lasting yet reversible changes in behavioral state remain poorly understood. This gap is particularly pronounced for complex behaviors such as seasonal migration, where the molecular and neural pathways linking seasonal environmental inputs to directed movements are only beginning to be elucidated.

Monarch butterflies (*Danaus plexippus*) offer a particularly compelling model system to explore this problem. Each fall, eastern North American monarchs undergo a seasonal long-distance migration from their breeding grounds in the northern United States and southern Canada to their overwintering sites in the central part of the Mexican Transverse Volcanic System^1–4^. Fall migrants travel thousands of kilometers southward in a state of reproductive quiescence, overwinter in this arrested state, before reproduction is initiated in the spring and mated migrants move northward to initiate the first leg of their return journey^1,3,5^. Flight orientation during both southward and northward migrations is primarily guided by a bidirectional, time-compensated sun compass^6–9^. Directional skylight cues are detected by the eyes and integrated within the central complex, a midline structure of the brain that serves as the insect navigation center^10,11^. This compass information is time-compensated by circadian clocks in the antennae, which adjust for the changing solar position throughout the day^12,13^.

Intriguingly, prolonged exposure to cold has been identified as the environmental cue that seasonally reverses flight orientation from south in the fall to north in the spring (Fig. 1a)^8^. Fall migrants experimentally exposed to cold temperatures mimicking overwintering conditions for as little as four weeks exhibit northward flight, recapitulating the natural flight orientation of spring remigrants. In contrast, aged fall migrants maintained in warmer, fall-like conditions continue to orient south^8^. Despite this striking behavioral plasticity, the molecular and cellular mechanisms by which cold exposure induces this reversal in flight orientation, and thereby ensures the completion of the migratory cycle, remain unexplored.

**Figure 1:**
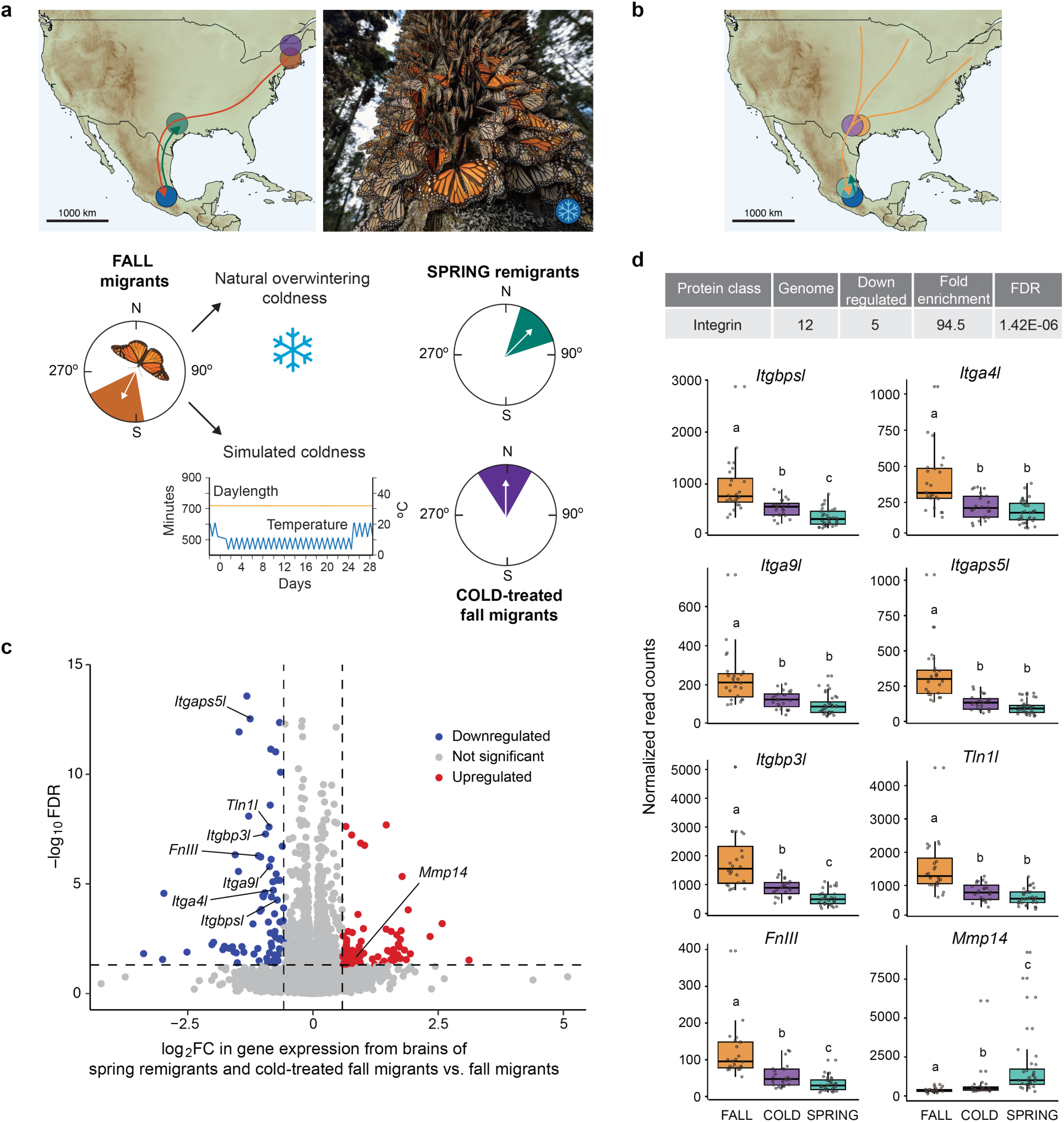
Cold exposure experienced by overwintering monarch butterflies elicits downregulation of integrin signaling in the brain. **a**, Representation of the effect of natural and simulated overwintering coldness on orientation behavior of Eastern North American migratory monarchs. Monarchs caught and tested in the fall (Massachusetts, MA; orange circle) orient south/southwest, and those caught and tested in the spring (Texas; green circle) orient north/northeast. Monarchs caught in the fall in MA and exposed to 24 days of premature cold exposure mimicking overwintering coldness (cold-treated; purple circle) reverse their flight orientation north. Modified from *Guerra and Reppert 2014*. Picture shows monarchs at their overwintering sites in Mexico (blue circle; photo credit: Jaime Rojo). **b**, Monarchs used in this study were captured in the fall in Texas and exposed to fall conditions (orange circle) or cold exposure (purple circle) as in *Guerra and Reppert 2014*. Spring remigrants were captured at the overwintering sites at the time of their departure northward (green circle). **c,** Volcano plot depicting differentially expressed genes in brains of spring remigrants/cold-treated fall migrants relative to fall migrants. FC, fold change; FDR, false discovery rate. Dotted lines, threshold of statistical significance. **d**, *Top*, Functional enrichment analysis reveals integrin as a protein class significantly downregulated in brains of spring remigrants and cold-treated fall migrants. *Bottom*, Box plots and raw data showing normalized read counts of expression for the five integrin encoding genes and genes associated with integrin-mediated processes in brains of fall migrants, cold-treated fall migrants and spring remigrants (n = 24 per condition). Error bars on box plots represent 1.5 times the interquartile range. Statistics were obtained by two-tailed Kruskal-Wallis test and post-hoc pairwise comparisons using two-tailed Dunn test with Bonferroni correction. Statistically significant differences between conditions are indicated by different letters (*p* < 0.05). *Itgbpsl*, *integrin beta PS-like* (Kruskal–Wallis χ² = 38.6, *p* = 3.95e^−9^); *Itga4l*, *integrin alpha 4-like* (χ² = 24.9, *p* = 3.79e^−6^); *Itga9l*, *integrin alpha 9-like* (χ² = 31.1, *p* = 1.75e^−7^); *Itgaps5l*, *integrin alpha PS5-like* (χ² = 43.7, *p* = 3.17e^−10^); *Itgbp3l*, *integrin beta pat3-like* (χ² = 44.5, *p* = 2.14e^−10^); *Tln1l*, *talin-1-like* (χ² = 35.7, *p* = 1.72e^−10^); *FnIII*, *fibronectin type III (*χ² = 43.28, *p* = 4.00e^−10^); *Mmp14*, *matrix metalloproteinase 14* (χ² = 42.4, *p* = 5.98e^−10^).

Here, we investigate the molecular and cellular basis of this cold-induced behavioral reprogramming. Using bulk and single-cell transcriptomic analyses, we show that exposure to cold is accompanied by coordinated transcriptional changes in the brain, prominently involving genes associated with extracellular matrix (ECM) signaling. In particular, *integrins*, encoding key ECM receptors, are consistently downregulated in the brains of both cold-exposed fall migrants and natural spring remigrants. Building on previous evidence of divergent selection in collagen type IV alpha 1 subunit, a major ECM component, between migratory and non-migratory populations, our data point to dynamic remodeling of ECM signaling as a potential regulator of seasonal flight orientation. Cellular mapping further suggests that cold exposure alters interactions between integrins in surface glia and ECM deposited by hemocytes, which together form the blood-brain barrier (BBB), and physiological evidence suggests it modulates BBB permeability. These findings identify the BBB as a site of environmentally driven plasticity and suggest that seasonal modulation of BBB properties may contribute to migratory state transitions in monarch butterflies.

## Results

### Cold exposure reprograms the rhythmic transcriptome in monarch brains

Sustained environmental exposure plays a key role in reshaping behavioral outputs across seasons. This suggests that environmentally driven regulation may reprogram brain gene transcription, enabling flexible, season– and context-dependent behavioral plasticity^14–16^. To investigate this possibility in the context of cold-induced reversal of flight orientation in monarch spring remigrants, we started our investigation by profiling and comparing the transcriptomes in the brains of wild-caught fall migrants maintained under fall-like conditions (fall migrants) with those exposed to 24 days of simulated overwintering-like coldness followed by 4 days of fall-like temperatures (cold-treated fall migrants; Fig. 1b), as previously described^8^. As a natural comparison, we also analyzed brains of wild-caught spring remigrants captured at departure from their overwintering sites in the spring, after natural exposure to the full winter cold period (Fig. 1b). Because migratory monarchs use a bidirectional sun compass that is time-compensated by circadian clocks^8^, we reasoned that cold exposure might alter compass output in the brain in a time-of-day-dependent manner. Accordingly, we profiled the transcriptomes of fall migrants, cold-treated fall migrants and spring remigrants at 3-hour intervals throughout the 24-hour day using RNA-seq (Supplementary Tables 1-3).

We first assessed whether cold exposure affected the rhythmic expression of core clock components, as changes in flight direction could arise from temperature-dependent modulation of circadian gene rhythmicity. A differential rhythmicity analysis using *dryR*^17^ identified 1,996 rhythmic genes expressed in phase in the brains of all seasonal forms (Supplementary Fig. 1a; Supplementary Table 4). Among these were the core circadian clock genes *period*, *timeless* and *cryptochrome 2*, which are known to exhibit 24-hour cycling in the brains of wild-type monarchs raised in summer conditions^18^ (Supplementary Fig. 1b), suggesting that cold exposure does not disrupt core brain clock function.

We next tested whether rhythmic gene expression could be differentially regulated by cold exposure acting in synergy with the circadian clock, as previously shown in other physiological contexts^19,20^. Although differential rhythmic gene expression was observed in several pairwise comparisons between fall migrants, cold-treated fall migrants and spring remigrants (Supplementary Fig. 1c, d, 2a-d and Supplementary Tables 4-16), we focused on genes that were similarly regulated in the brains of cold-treated fall migrants and spring remigrants but differentially regulated in fall migrants. We did not detect any genes whose rhythmicity was in anti-phase between groups, which could have potentially explained the seasonal reversal in flight orientation. However, we identified 207 genes that were rhythmic in the brains of fall migrants but arrhythmic in cold-treated fall migrants and spring remigrants (Supplementary Fig. 1c and Supplementary Table 5), and 363 genes that were arrhythmic in fall migrants but rhythmically expressed in phase in cold-treated fall migrants and spring remigrants (Supplementary Fig. 1d and Supplementary Table 6). No significant enrichment for biological processes or protein classes was found in either dataset, making it difficult to predict if any of these genes could contribute to flight orientation reversal in migratory monarchs.

**Figure 2:**
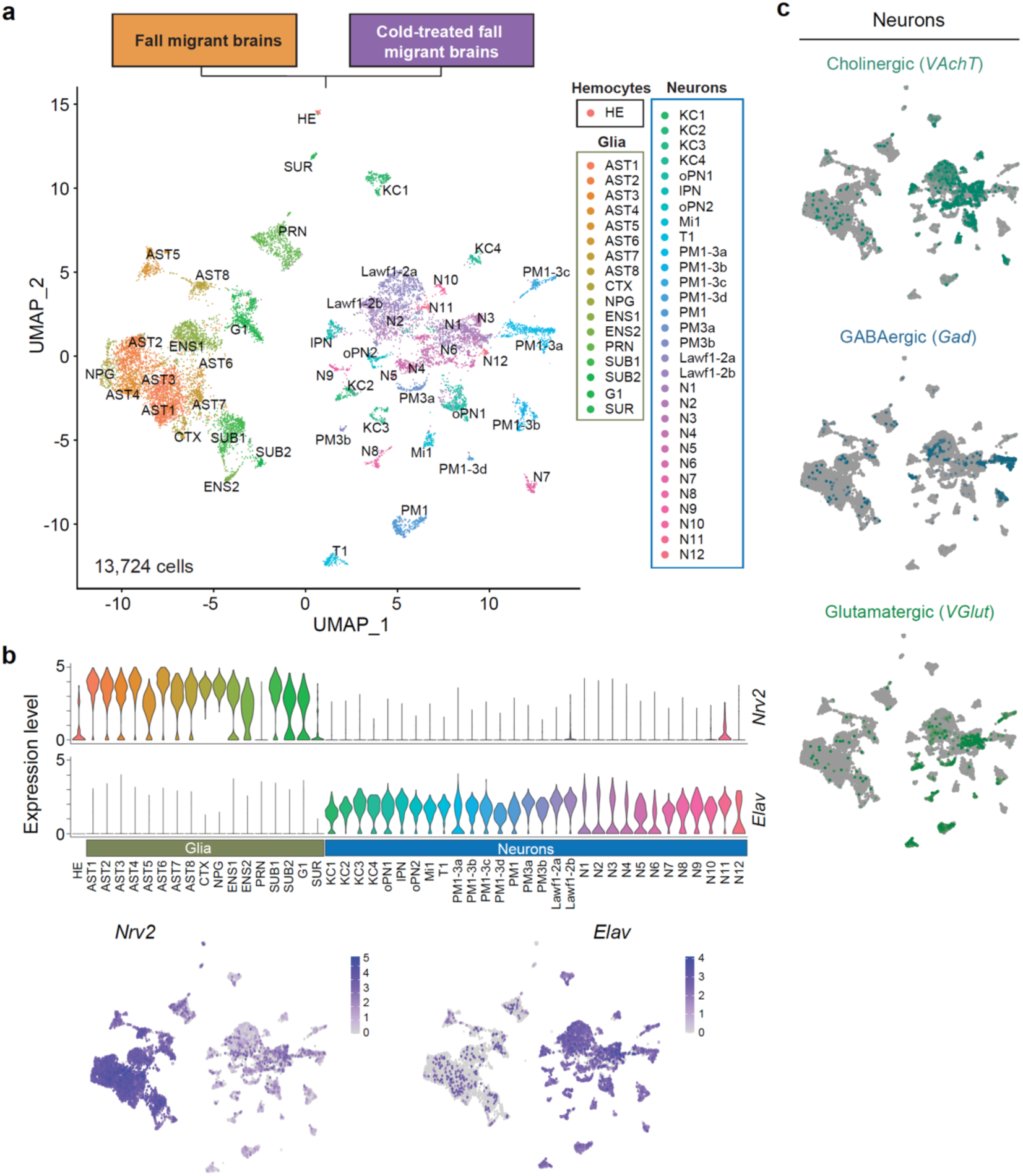
Single-cell transcriptome profiling of brains of adult fall migratory monarch butterflies and cold-treated fall migrants. **a**, Annotated UMAP visualization of integrated fall migrants and cold-treated fall migrants (3 pooled brains in duplicates for each) scRNA-seq data from >13K cells. Clusters were manually curated for expression of known marker genes and annotated (HE, hemocytes; AST, astrocyte-like glia; CTX, cortex glia; NPG, neuropil glia; ENS, ensheathing glia; PRN, perineurial glia; G1, unannotated glial cluster; SUR; surface glia; KC, Kenyon cells; oPN, olfactory projection neurons; IPN, insulin producing neuron; Mi, neurons of the optic lobe; T1, neurons of the optic lobe; PM, neurons of the optic lobe; Lawf, lamina wild-field neurons; N1-12, unannotated neuronal clusters. **b,** *Top*, Example of selected marker genes for cluster annotation presented as expression level per cluster; graphs depicts glial (*Nrv2*) and neuronal (*Elav*) marker genes. *Bottom*, Featureplots of marker genes showing per-cell expression levels. **c,** Featureplots of neuronal subtypes classification based on per-cell expression of major neurotransmitters. *VAChT* indicates cholinergic neurons, *Gad* indicates GABAergic neurons, and *VGlut* indicates glutamatergic neurons.

Nevertheless, these data demonstrate a role for seasonal cold exposure in reprogramming the rhythmic transcriptome in monarch brains. Two notable examples of this cold-induced reprogramming involve the core circadian activator *Bmal1* and an *octopamine receptor*. *Bmal1*, previously reported to be arrhythmic in the brains of summer monarchs and fall migrants^18,21^, exhibited robust rhythmicity in the brains of both cold-treated fall migrants and spring remigrants, suggesting cold-dependent rhythmic transcriptional or post-transcriptional regulation (Supplementary Fig. 1e). In addition, a gene encoding a receptor for octopamine, a potent neuromodulator of behavior in insects^22^, was arrhythmic in fall migrants but rhythmic and upregulated by ∼50-fold in the brains of monarchs exposed to cold (Supplementary Table 6), implicating neuromodulatory pathways in the seasonal plasticity of migratory behavior.

### Cold exposure attenuates integrin signaling and promotes ECM remodeling in monarch brains

Because cold-induced flight orientation reversal may occur independently of the circadian clock’s time compensation mechanism, we next examined differential gene expression in the brain in a time-of-day-independent manner. To do so, we pooled all samples for each seasonal form and compared global transcriptomic profiles using all samples as replicates. We identified 161 upregulated and 103 downregulated genes in the brains of cold-treated fall migrants relative to fall migrants (Supplementary Tables 17-18). Likewise, 316 genes were upregulated and 306 genes were downregulated in spring remigrants compared with fall migrants (Supplementary Tables 19-20), an overall increase likely reflecting that spring remigrants experience additional environmental factors in the wild beyond cold exposure alone.

Intersecting the two datasets revealed 65 genes consistently upregulated and 65 genes consistently downregulated in the brains of monarchs exposed to cold (Fig. 1c and Supplementary Table 21). Although enrichment analysis did not identify biological processes significantly enriched among upregulated genes, three of the most highly upregulated genes out of the 65 encoded glycine-rich RNA-binding proteins (GRPs; Supplementary Fig. 3a, b). GRPs are known to increase in abundance under cold conditions and contribute to cold tolerance in plants^23^. Given the parallel between overwintering cold exposure in monarchs and vernalization in plants, these GRPs could represent unexplored molecular substrates for cold tolerance in insects.

In contrast, enrichment analysis of downregulated genes identified *integrins* as significantly over-represented, with a fold-enrichment of 94.50 (Fig. 1d). Of the twelve *integrin* genes present in the monarch genome, five, annotated as *integrin beta PS-like*, *integrin alpha 4-like*, *integrin alpha 9-like*, *integrin alpha PS5-like*, and *integrin beta pat3-like* were significantly downregulated in the brains of both cold-treated fall migrants and spring remigrants (Fig. 1c, d). In the brain, integrins are broadly expressed in both neuronal and glial cell types and are known to regulate neuronal connectivity and synaptic plasticity^24^. Acting as heterodimeric cell-surface receptors composed of alpha and beta subunits, they transduce extracellular signals by bridging the extracellular matrix (ECM), a macromolecular network of collagens, fibronectins, elastins, laminins, and glycoproteins, to the intracellular cytoskeleton^25,26^.

Interestingly, a population genomics study comparing migratory and non-migratory monarch populations previously identified *collagen type IV alpha 1*, a key ECM component, as one of three genes present within a 21-kb genomic region associated with migratory behavior and under strong divergent selection^27^. In this context, our finding raised the intriguing possibility that *collagen type IV alpha 1* and the *integrin* genes may functionally interact in the brain. To investigate this idea further, we mined our list of differentially expressed genes for additional candidates associated with integrin-mediated processes. Among downregulated genes, we identified *talin-1-like*, which encodes an adaptor protein linking integrins to the actin cytoskeleton^25^, and LOC116772305, an unannotated gene containing a domain of fibronectin type III, an ECM glycoprotein involved in matrix assembly^28^, which we hereafter call *fibronectin type III* (Fig. 1c, d). Conversely, *matrix metalloproteinase-14*, which encodes an enzyme that degrades ECM components^29^, was significantly upregulated in the brains of cold-treated fall migrants and spring remigrants (Fig. 1c, d).

Taken together, these data suggest that sustained cold exposure, which drives the reversal of flight orientation in spring migratory monarchs^8^, attenuates integrin signaling and also induces ECM remodeling in the brain, potentially mediating neuronal and/or glial plasticity underlying seasonal behavioral transitions.

### scRNA-seq identifies discrete neuronal and glial cell types in the monarch brain

Given the intimate association of integrins/ECM remodeling with a number of neurobiological processes in the brain, including synaptic transmission, scaling and plasticity^25^, we next sought to define in which cell types the downregulated *integrins* were expressed. We performed single-cell RNA sequencing (scRNA-seq) on dissociated adult brains (containing optic lobes) from fall migrants and cold-treated fall migrants. After filtering, integration and cell clustering using Seurat^30^, we obtained 13,724 high-quality brain cells that grouped into 48 cell clusters (Fig. 2a). Cell type identity was inferred using cluster-specific marker genes characterized in *Drosophila melanogaster*^31–33^.

Neuronal and glial cell populations were respectively identified based on the expression of *elav* and *nervana2* (Fig. 2b), and further separated into more discrete cell types using additional markers (Supplementary Fig. 4, 5 and Table 22). Among neuronal cell types, we identified the three major neurotransmitter-defined classes, cholinergic, GABAergic, and glutamatergic using their respective known marker genes, *vesicular acetylcholine transporter* (*VAchT*), *glutamic acid decarboxylase* (*Gad*), and *vesicular glutamate transporter* (*VGlut*)^32,33^ (Fig. 2c). Of the 30 neuronal clusters, we were able to annotate 18 to specific subtypes, including Kenyon cells of the mushroom body (*Vmat*)^32,33^, olfactory projection neurons (*acj6* and *vvl*)^33^, insulin-producing neurons (*Tk*), neurons of the optic lobes (*hth* and *bsh* for M1^31,32^, *eaat1* for T1^32^ and *lim 3* for PM^31,32^) as well as lamina wide-field neurons (*eya*)^32^ (Supplementary Fig. 4 and Table 22). We also identified 17 discrete glial clusters, including the following major glial classes: astrocyte-like glia (*Eaat1, Gs2*)^31^, cortex glia (*wrapper*)^31,32^, neuropil glia (*ebony*)^31^, ensheathing glia (*draper*, *Gs2*)^31^, and surface glia (*indy*)^32^ (Supplementary Fig. 5 and Table 22). Although limited in scope in the context of this study, this analysis establishes the first single-cell atlas of the monarch brain, providing a framework for examining cell-type specific expression changes associated with cold exposure.

### *Integrins* and transcripts associated with integrin-mediated processes and ECM remodeling localize to surface glia of the blood-brain barrier

To determine the cellular origin of the integrin and ECM remodeling signatures identified in our bulk RNA-seq analysis, we examined the single-cell expression patterns of *integrin* subunits and key associated transcripts from integrated scRNA-seq data of fall migrants and cold-treated fall migrants. Strikingly, all were predominantly expressed within a single cell cluster (Fig. 3a). The molecular profile of this cluster, characterized by enriched expression of *Indy*, a conserved surface glia marker in other insects^32,34^, indicates that integrins and their associated ECM components localize to surface glia (Fig. 3a, b).

**Figure 3:**
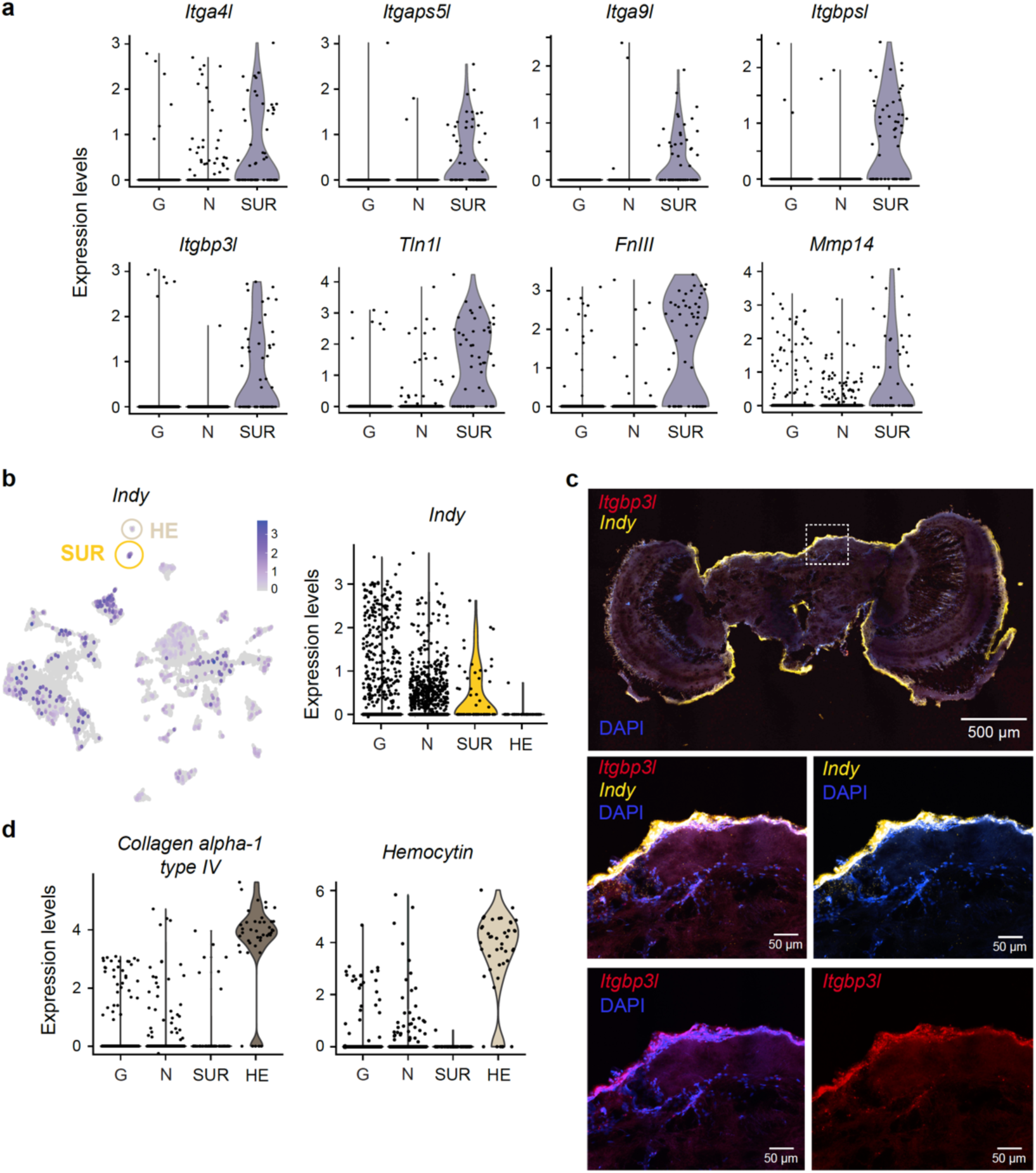
Surface glia specific expression of *integrins* and *integrin*s-associated genes differentially expressed in brains exposed to overwintering-like coldness. **a**, Violin plots of *Itgbpsl*, *Itga4l*, *Itga9l*, *Itgaps5l*, *Itgbp3l*, *Tln1l*, *FnIII*, and *Mmp14* expression levels showing specific expression in surface glia (SUR) as compared to other glial (G) and neuronal (N) cell types. **b,** Featureplot of the homologue of *Drosophila* surface glia marker *Indy* in monarch brains (*left*) and violin plots of its expression levels showing significant enrichment in surface glia compared to other glial, neuronal and hemocyte (HE) cell types (*right*). **c,** Immunofluorescence of *Itgbp3l* and *Indy* in the monarch brain counterstained with 4′,6-diamidino-2-phenylindole (DAPI). Top image shows the peripheral localization of *Itgbp3l* and *Indy* on a 25 µm brain cryosection, and bottom four images show magnifications of the inset depicting the co-localization of *Itgbp3l* and *Indy*. **d,** Violin plots of *Collagen alpha 1 type IV* and *Hemocytin* expression levels showing specific expression in hemocytes (HE).

Surface glia constitute the outermost layer of the insect blood-brain barrier (BBB), which separates the nervous system from the hemolymph of the open circulatory system^35^. This layer comprises two morphologically distinct glial types: apical perineurial cells, which are covered by the ECM-rich neural lamella, and basal subperineurial glia^36^. Although we could not molecularly identify perineurial glia in our single-cell dataset, none of the *integrin* genes were expressed in subperineurial glia (identified by *moody* expression; Supplementary Fig. 5 and Table 22), strongly suggesting their expression to be restricted to apical perineurial cells, in line with their function as ECM receptors. To confirm the anatomical localization of the *integrin* subunits to the BBB, we performed fluorescent *in situ* hybridization using RNAScope against one of the integrins, *Itgbp3l*, on monarch brain cryosections. As predicted, *Itgbp3l* co-localized with *indy* in a layer of cells encasing the brain neuropiles (Fig. 3c). Consistent with results from bulk RNA-seq in brains, 3 of the 5 *integrins*, together with *Talin 1-like* and *Fibronectin type III*, showed a significant decrease in expression at the single-cell level in surface glia of cold-treated fall migrants relative to fall migrants (Supplementary Fig. 6a) based on an overall reduction in the number of cells which expressed them (Supplementary Fig. 6b).

Finally, to assess whether *collagen type IV alpha 1*, previously identified as a candidate gene for migratory behavior^27^ may be associated with BBB function, we investigated its cellular source. Because collagen type IV is known to be secreted by hemocytes, insect blood cells circulating in the hemolymph, and incorporated into the ECM-rich neural lamella^37,38^, we hypothesized that this gene might be expressed in these cells. We indeed found that *collagen type IV alpha 1* was predominantly expressed in a distinct hemocytic cell cluster based on the specific expression of the *hemocytin* marker (Fig. 3d). Interestingly, *matrix metalloproteinase 14*, an enzyme that degrades ECM components^39^ and predominantly expressed in surface glia (Fig. 3a), was upregulated in brains of monarchs exposed to cold (Fig. 1c, d), suggesting that in addition to attenuating integrin signaling at the BBB, cold exposure concurrently affects ECM integrity.

Together, these findings strongly suggest that cold exposure remodels the molecular and cellular architecture of the BBB, establishing it as a dynamic interface through which environmental temperature may drive seasonal reprogramming of migratory behavior.

### Cold exposure functionally alters monarch BBB permeability

Given that integrin signaling and ECM components are critical for maintaining the structural integrity of the BBB, we hypothesized that their cold-induced downregulation may compromise barrier function and allow greater penetration of circulating factors into the brain. To test this prediction, we assessed BBB permeability *in vivo* in laboratory-raised monarchs maintained in fall conditions or exposed to 26 days of cold, using a fluorescent tracer diffusion assay commonly applied in invertebrate models^40,41^ (refs; Fig. 4a). Monarchs were injected at midday in the thorax with fluorescein, a classical 332 Da BBB tracer, and brains and hemolymph were collected at midday for quantification. Fluorescein levels in the brain were ∼2-fold higher in brains of cold-exposed monarchs relative to fall controls (Fig. 4b), demonstrating that cold exposure functionally increases BBB permeability.

**Figure 4:**
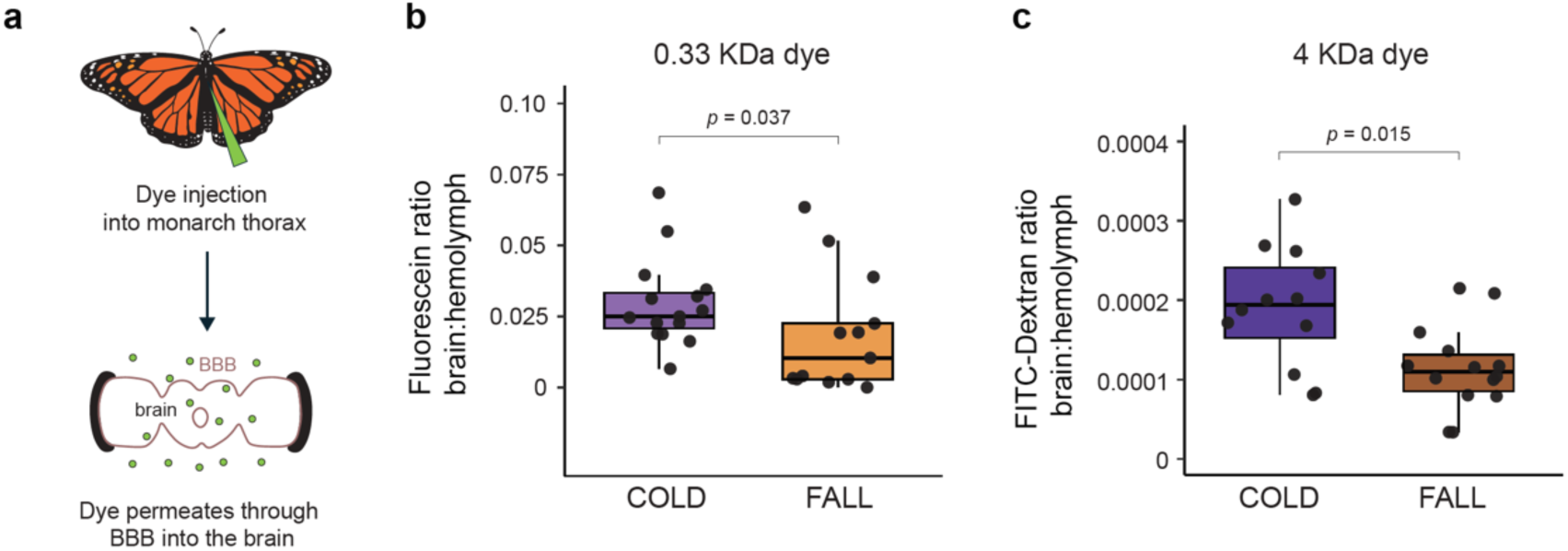
Cold-induced increased permeability of the blood-brain barrier (BBB). **a**, Workflow illustrating the fluorescent tracer diffusion assay to assess BBB permeability. **b**, Quantification of fluorescein ratios of brains: hemolymph of monarchs subjected to cold treatment (n = 15) or maintained in fall-like (n = 13) conditions (two-tailed Mann-Whitney U test; w = 143, *p* = 0.037). **c**, Quantification of FITC-dextran ratios of brains: hemolymph of monarchs subjected to the same conditions (cold n = 12; fall n =14) as in b (two-tailed Wilcoxon Mann-Whitney; w = 131, *p* = 0.015). In **b-c** dots represent individual butterfly marker levels, boxes represent the 25^th^ to 75^th^ percentile, the bar across the middle represents the median and error bars represent 1.5x IQR.

Because this increase appears to be passive, consistent with lack of evidence for differential expression of known active transporter genes between fall and cold exposed monarchs, we next asked if larger molecules could also penetrate the BBB under cold conditions. Repeating the assay with a 4 KDa fluorescein isothiocyanate (FITC)-dextran revealed significantly higher FITC-dextran accumulation in the brains of cold-exposed monarchs relative to fall controls (Fig. 4c), albeit at overall concentration lower than fluorescein.

Together, these findings indicate that seasonal cold exposure weakens the structural and functional integrity of the monarch BBB, enhancing its permeability to circulating molecules in the hemolymph that could facilitate temperature-dependent molecular signaling between the periphery and the brain to recalibrate migratory flight orientation.

## Discussion

Animal seasonal migration represents a striking example of behavioral plasticity, with individuals reversing migratory orientation seasonally. While the sensory and circadian mechanisms guiding migratory orientations are well characterized^1,15,42,43^, the molecular and cellular underpinnings of migratory behavior and their seasonal transitions have remained largely unresolved. Using Eastern North American monarch butterflies as a model, we show that exposure to cold, the environmental cue experienced during overwintering that reverses flight orientation from south to north^8^, elicits coordinated transcriptional and physiological changes in the brain. Cold exposure reprograms the brain transcriptome and attenuates integrin-mediated extracellular matrix (ECM) signaling in surface glia at the blood-brain barrier (BBB), coinciding with increased BBB permeability. To our knowledge, this is the first demonstration that seasonal environmental cues induce transcriptional and functional changes in the BBB, suggesting it likely acts as a dynamic interface through which seasonal signals may influence neural circuits underlying migratory behavior.

Integrins function as heterodimeric receptors linking ECM proteins such as collagen, fibronectin and laminins, to the cytoskeleton via adaptor proteins such as talin^25,44^. The consistent and coordinated downregulation of *integrin* subunits, *talin*, and *fibronectin type III* in both experimentally cold-treated and naturally remigrating monarchs, together with upregulation of an enzyme capable of degrading all kinds of ECM proteins, *matrix metalloproteinase 14* (*Mmp14*), indicates that cold exposure weakens integrin-ECM interactions and may promote ECM turnover. In combination with prior evidence of divergent selection on *collagen type IV alpha 1* between migratory and non-migratory monarch populations^27^, these results support the ECM-integrin axis as a molecular substrate for the seasonal plasticity of migratory orientation behavior.

Unexpectedly, single-cell transcriptomics and *in situ* mapping localize the cold-responsive integrin signature to surface glia. Surface glia comprise the outer perineurial glia and the inner subperineurial glia and form the insect BBB together with the neural lamella composed of a dense network of ECM proteins. Although our dataset did not differentiate perineurial from subperineurial glia based on the expression of known specific markers in *Drosophila*, the cell cluster in which integrin genes are expressed is clearly separated from subperineurial glia and expressed the surface glia marker *indy*, supporting its perineurial nature. By weakening perineurial glia-ECM interactions, cold exposure may reduce the barrier function of the perineurial layer, as evidenced by increased passive diffusion of tracers into the brain.

Together, our findings indicate that the BBB is a dynamic structure capable of responding to environmental cues to possibly regulate seasonal behavior, adding to its known dynamic regulation by circadian rhythms and sleep in fruit flies and mammals^45,46^. Direct genetic demonstration that cold-induced BBB permeability caused by integrin downregulation is responsible for the switch in migratory behavior is currently limited by technical constraints, including inefficiency of RNA interference knockdown in lepidopterans and unknown conditions necessary to fully induce the migratory phenotype from laboratory-raised CRISPR mutants.

Nevertheless, seasonal modulation of BBB permeability could enable peripheral signals to influence the activity of compass circuits in the central complex, a brain structure known as the navigation center in insects, potentially contributing to seasonal behavioral plasticity. Increased permeability caused by weakened integrin-ECM interactions after cold exposure may allow endocrine or metabolic factors that fluctuate in the hemolymph with overwintering conditions to permeate through the brain to influence neuronal activity in the central complex (Fig. 5). This model would provide a parsimonious explanation for how prolonged environmental cold exposure could cause a switch in migratory orientation without requiring the formation of new neural pathways. Trafficking of molecules from the periphery into the brain through the BBB to regulate behavior is not without precedent, as active transport of lipids through the *Drosophila* BBB has recently been shown to modulate sleep^47,48^. Future work identifying the specific factors that cross the monarch BBB after cold exposure and the neurons they affect will be critical for establishing a causal link between seasonal BBB remodeling and changes in orientation behavior.

**Figure 5:**
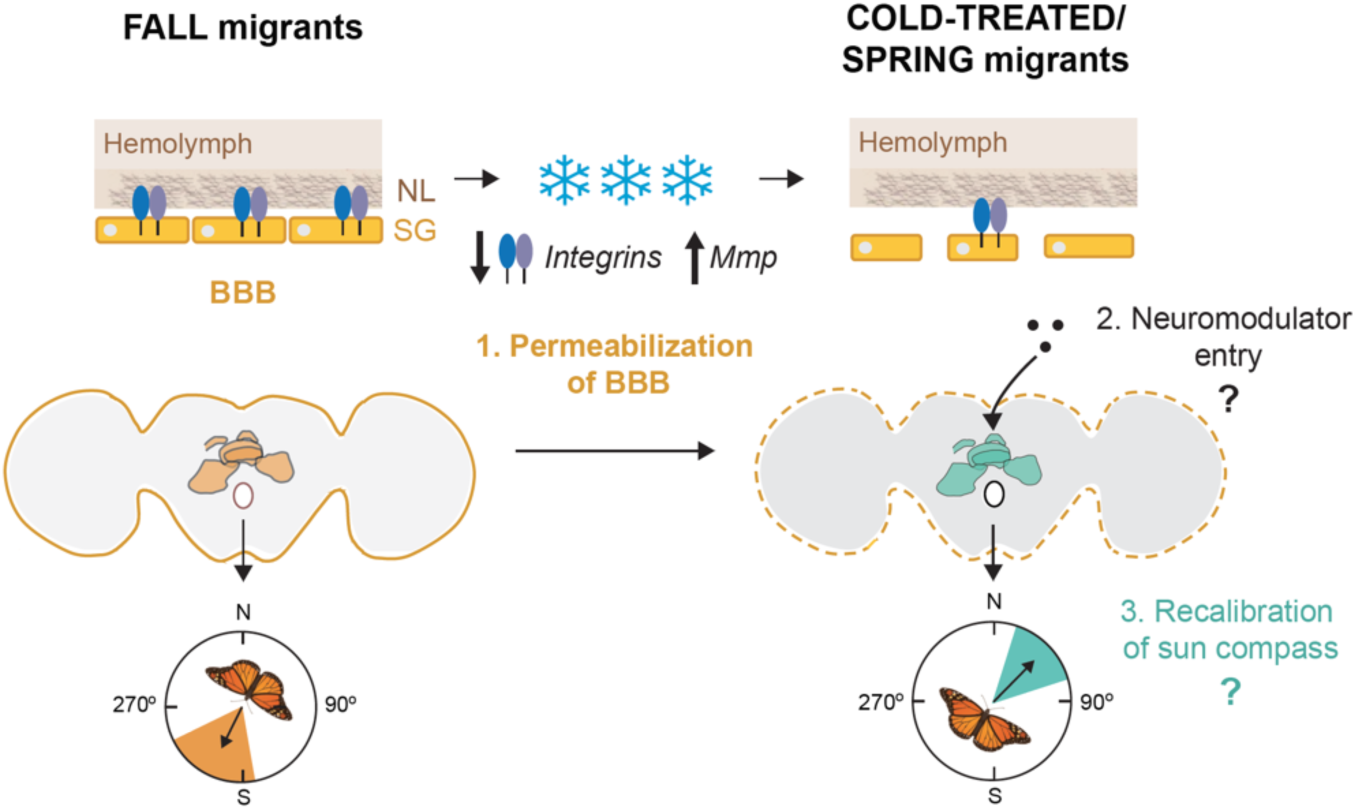
Model for cold-induced modulation of the BBB underlying seasonal behavioral reprogramming in monarchs. Downregulation *of integrins* and genes encoding intracellular and extracellular binding partners (e.g. *talin* and *fibronectin type III*) and upregulation of genes encoding matrix metalloproteinases degrading tight junctions in surface glia upon sustained exposure to overwintering-like coldness results in increased permeabilization of the BBB. BBB permeabilization could in turn lead to flight orientation reversal by allowing the entry of circulating molecules from the periphery into the brain capable of recalibrating the brain sun compass neural circuitry. The colored structure in the monarch brain represents the central complex, where orientation relative to cues and goal direction is encoded.

More broadly, our findings highlight a potentially conserved role for the BBB in mediating environmentally induced behavioral plasticity. In both fruit flies and mammals, circadian modulation of barrier permeability has been shown to influence sleep, neural function and metabolic homeostasis^41,47,48^. Our results extend this concept to seasonal timescales, suggesting that dynamic regulation of the BBB may represent a general strategy for coordinating physiological and behavioral responses to changing environments. Comparative studies in other migratory or hibernating species could help determine whether modulation of barrier permeability is a common mechanism underlying seasonal adaptation. Finally, these findings highlight a potential vulnerability of monarch migration to climate change, as rising temperatures could disrupt BBB-mediated behavioral transitions.

## Methods

### Monarch butterflies sources

Monarch butterflies used for sequencing experiments were captured during their fall migration in College Station, TX (30°37’N, 96°20’W) and at their overwintering sites at Sierra Chincua, Michoacán, Mexico (19°37’N, 100°17’W) just prior to their spring northward remigration. Spring remigrants were captured with permit SGPA/DGVS/12334/19 to M.I. Ramírez. Monarch butterflies used for immunohistochemistry and blood-brain barrier permeability assessment were raised in the laboratory on semi-artificial diet from eggs to adults under 15-hour light:9-hour dark cycles at 25°C, as previously described^49,50^.

### Bulk RNA-sequencing

Fall migrants were captured on October 23-24, 2017 in College Station, TX (30°37’N, 96°20’W). One group was subjected to 12 hours of light at 11°C and 12 hours of dark at 4°C to simulate overwintering coldness for 24 days before being switched to warmer conditions of 12 hours of light 21°C and 12 hours of dark at 12°C for 4 days, as previously described (thereafter called cold-treated fall migrants)^8^. Another group was maintained under fall-like conditions of 11 hours of light at 21°C and 13 hours of dark at 12°C for 30 days (fall migrants). Wild-caught spring remigrants that experienced natural overwintering coldness were collected just prior to their expected northward departure on March 3, 2020 at Sierra Chincua, Mexico (19°37’N, 100°17’W). All animals were kept in glassine envelopes and manually fed every 2 or 3 days with 25% honey solution, except for spring remigrants which were fed only once after their capture before dissections.

At the end of the 30 days of treatment, three pooled brains of fall migrants and of cold-treated fall migrants were dissected in triplicates at 3-hour intervals throughout a 24-hour day starting from Zeitgeber 0 (ZT0; light on) to ZT 21 in 0.5X RNAlater, snap frozen in liquid nitrogen and stored at −80°C. Similarly, four replicates of three pooled brains of spring remigrants were dissected the day following their capture, stored in 1X RNAlater at 4°C, transported in frozen gel ice packs, and stored at −80°C upon return to the lab. All dissections during the dark phase of the light:dark cycle were performed under red light.

Total RNA was extracted using RNeasy Mini kit (Qiagen) and quality control was performed on an Agilent tape station. PolyA+ RNA was isolated from 750 ng of total RNA with NEBNext Oligo d(T) magnetic beads, and libraries were prepared using the NEBNext Ultra II Directional RNA Library Prep kit and NEBNext Multiplex Oligos, following the manufacturer’s recommendations. After verification of library quality and expected insert size using a tape station, equimolar concentrations of libraries were pooled from samples of fall migrants and cold-treated fall migrants, as well as from samples of spring remigrants for multiplexing. The pooled samples were sequenced on two lanes of NovaSeq 6000 with paired end read lengths of 150 bp at the North Texas Genome Center, a partner of the Texas A&M Institute of Genome Sciences and Society. Demultiplexed raw data were pre-processed using BBDuk to obtain high quality reads, which were mapped to the chromosome-level monarch genome^51^ using STAR^52^. After the first pass of mapping, the spliced junctions generated from each sample were combined and used for a second pass mapping generating more accurate alignments. Transcript abundances were then estimated using Salmon^53^ and a count matrix containing gene-level expression values were generated using the R package *tximport*^54^. Genes with no mapped reads in all samples were filtered out of the matrix and not considered in subsequent analysis.

### Identification of differentially expressed genes

To identify genes whose mRNA abundance oscillates throughout the course of the 24-hour day in the brains of fall migrants, cold-treated fall migrants, and spring remigrants, we used the R package *dryR*^17^ to simultaneously perform rhythmicity detection and differential rhythmicity analysis across the three conditions. Analysis of differential gene expression in a time-of-day-independent manner between fall migrants and cold-treated fall migrants or spring remigrants was performed using DESeq2^55^ considering all samples from each seasonal form as replicates irrespective of their time of collection. Genes were considered differentially expressed if adjusted *p*-value < 0.05 and log2 fold-change > 0.585 (*i.e.*, fold-change > 1.5) for up-regulated genes or < −0.585 (*i.e.*, fold-change < 0.667) for down-regulated genes.

### Enrichment analysis

Enrichment of biological processes and/or protein classes among rhythmic and differentially expressed genes was performed using the statistical overrepresentation test of PANTHER^56^, with all genes expressed in the brains of each condition used as background.

### Single-cell RNA-sequencing

Fall migrants for single-cell RNA-seq were captured on October 18-19, 2021, in College Station, TX (30°37’N, 96°20’W) and subjected either to fall-like conditions or overwintering-like coldness, similar to that described for bulk RNA-seq. Single-cell suspensions from brain tissues were prepared following a previously established protocol for *Drosophila* brain^32^. Briefly, three pooled brains were dissected in duplicates from fall migrants and cold-treated fall migrants between ZT5 and ZT6 in 1X DPBS, and immediately transferred to ice-cold 1X DPBS. After removal of the supernatant, each sample was mixed with 50 μl of 3 mg/ml dispase (MilliporeSigma), and 75 μl of 100 mg/ml collagenase I (ThermoFisher). Brains were dissociated using a thermoshaker at 25°C and 500 rpm for 2 hours. Every 15 minutes, the samples were mixed by gentle pipetting to facilitate enzymatic digestion of the extracellular matrix. Cells were washed with 1 ml of ice-cold 1X DPBS, resuspended in 500 μl of 1X DPBS supplemented with 0.04% BSA, and filtered through a 10-μm pluroStrainer (pluriSelect). Cell viability and concentration were determined using Moxi GO II (ORFLO). Individual cells were captured using 10X Genomics platform followed by cDNA synthesis, library preparations, and sequencing. Raw data were first processed using Cell Ranger^57^ for quality control, mapping to the chromosome-level monarch genome^51^, and quantification. Standard post-processing and clustering of cells were all performed using Seurat^30^, except for mitochondrial filtering. Prior to clustering, all data were integrated using Harmony^58^. Clusters of cells expressing mitochondrial genes were identified and removed from the data, which were re-processed, re-integrated, and re-clustered as described above.

### Immunohistochemistry

Brains from monarch butterflies held at 25°C under 15:9 hours light:dark conditions were dissected in Ringer’s solution, fixed for 24 hours in 4% paraformaldehyde and cryopreserved with increasing concentrations of sucrose at 10%, 20%, and 30% for approximately 18 hours each. Brains were mounted in Tissue-Tek (Sakura), frozen at −80°C overnight, and cryosectioned at 25 µm-thick slices with a cryostat.

*In-situ* hybridizations were performed using RNAscope Target Probes designed against specific candidates, a RNAscope Multiplex Fluorescent kit v2 (ACD) and TSA Vivid Dyes (ACD). To confirm the efficacy of the fixation and cryopreservation methods, we first used 1X RNAscope probes designed against the neuronal marker *elav* (targeting base pairs 6-1087 of *elav* mRNA XM_061528728.1) as a positive control, and a standard negative control probe designed against *Bacillus subtilis DapB* (https://acdbio.com/search/site/%252ADapB%252A/cms/probes/gene/dapb). TSA Vivid Fluorophore 520 was used at 1:750 to visualize the *elav* and *DapB* RNA bound probes. Cellular localization of *integrin beta pat-3-like* was assessed with a probe designed to target base pairs 113-1162 of its predicted mRNA (XM_032674126.1) and a probe targeting the surface glia marker *Indy* (base pairs 928-1878 of XM_032670464.2), both diluted to 1X in RNAscope Probe Diluent (ACD). Signal amplification and visualization was performed using TSA Vivid Fluorophore 650 for *integrin beta pat-3-like* and TSA Vivid Fluorophore 570 for *Indy*. Hybridization steps were performed according to the manufacturer’s instructions, with the exception of tissue preparation, for which slides were baked at 60°C for 1.5 hours. Following hybridization, slides were counterstained with 4′,6-diamidino-2-phenylindole (DAPI), mounted with ProLong Gold Antifade (Invitrogen) and imaged on a Nikon AX1R confocal microscope.

### Blood-brain barrier permeability

Laboratory-reared monarchs were subjected to either fall-like or overwintering-like conditions as described above for 26 days. Blood-brain barrier permeability assays were adapted from assays developed in flies^59^. Butterflies were injected manually between the thorax and the abdomen with a 1 ml syringe and 30 gauge (0.3 mm x 25 mm) needle (Becton, Dickinson and Company) containing 40 μL of a 1mM solution (0.37 mg/ml) of 0.33 KDa fluorescein (Sigma Aldrich) or a 2.5 mg/ml solution of 3-5 kDa FITC-dextran (Millipore Sigma) diluted in 1X PBS and blue food coloring to visually track circulation throughout the hemolymph. Injections were performed under a dissecting microscope between ZT4 and ZT8. Approximately 1-hour prior to injections, butterflies were fed 25% honey solution. Following injections of fluorescein, butterflies were held at 21°C for 10 minutes. Following injections of FITC-dextran, butterflies were held in their respective temperature treatments for 15 minutes. On each day of injections, two lab-reared butterflies were mock injected with a solution of 1X PBS and food coloring to account for background fluorescence. Brains were then rapidly dissected in 1X PBS under red light in darkness to prevent light-exposure of fluorescein/FITC and approximately 20 μL of hemolymph was collected from each butterfly. Brains were washed with 1X PBS and placed into 50 μL of 0.1% SDS in 1X PBS for at least 30 mins in darkness before being mechanically dissociated with a pestle. Collected hemolymph was supplemented with 30 μL of 1X PBS to a total of 50 μL. Fluorescein and FITC-dextran from brain and hemolymph samples (50 μL each) were measured at excitation/emission: 460/515 nm and 485/528 nm, respectively, using a Molecular Devices SpectraMax iD3 plate reader. For each set of measurements, a standard curve of 100%, 10%, and 1% of 1 mM fluorescein or 2.5mg/ml FITC-dextran respectively was used to calculate amount of dye in each sample. Fluorescence levels in the brain and hemolymph of all background butterflies were averaged, and these values were subtracted from each experimental butterfly’s brain and hemolymph fluorescence values to account for any tissue autofluorescence. Ratios of the fluorescence in the brain to hemolymph were then calculated for each butterfly.

### Quantification and statistical analysis

Details of statistical tests can be found in the figure legends. Kruskal-Wallis tests and post-hoc pairwise comparisons conducted using the Dunn test with Bonferroni correction were conducted using R Statistical Software version 4.1.1 and Mann-Whitney U-tests were conducted using R Statistical Software version 4.2.2.

### Data availability

Raw and processed files have been deposited to the Gene Expression Omnibus repository (https://www.ncbi.nlm.nih.gov/geo/query/acc.cgi?acc=GSE314314).

## Acknowledgements

We thank the Texas A&M Institute for Genome Sciences and Society for providing sequencing services, the Texas A&M High Performance Research Computing for providing computational resources and systems administration support; An Nguyen, Karen Ibarra for assistance with monarch maintenance and Samantha Iiams for assistance with reproductive status assessment of the dissected butterflies; Lamba Sangare and Jeff Jones for use of the SpectraMax iD3 plate reader and cryostat; Jaime Rojo for the picture of overwintering monarchs; and Paul Hardin and Steven Reppert for comments on the manuscript. The work was supported by IOS-1754725 grant from the National Science Foundation to C.M. and a Life Sciences Research Foundation/Welch Foundation postdoctoral fellowship to K.M.G.

## Author Contributions

A.B.L. and C.M. conceived the project. A.B.L. performed RNA-seq and scRNA-seq data collection and data analysis with the help of C.M., V.S. and Y.Z. K.M.G. performed *in situ* hybridizations and permeability assays and data analysis. M.I.R. and A.G.R. contributed to monarch specimens’ collection and access to infrastructure. A.B.L., K.M.G. and C.M. designed experiments and wrote the manuscript with input from all authors. All authors contributed to the discussion of the data and approved the final version of the manuscript.

