## Supplementary Information for "Seasonal blood-brain barrier plasticity links environmental cues to migratory behavior in monarch butterflies"

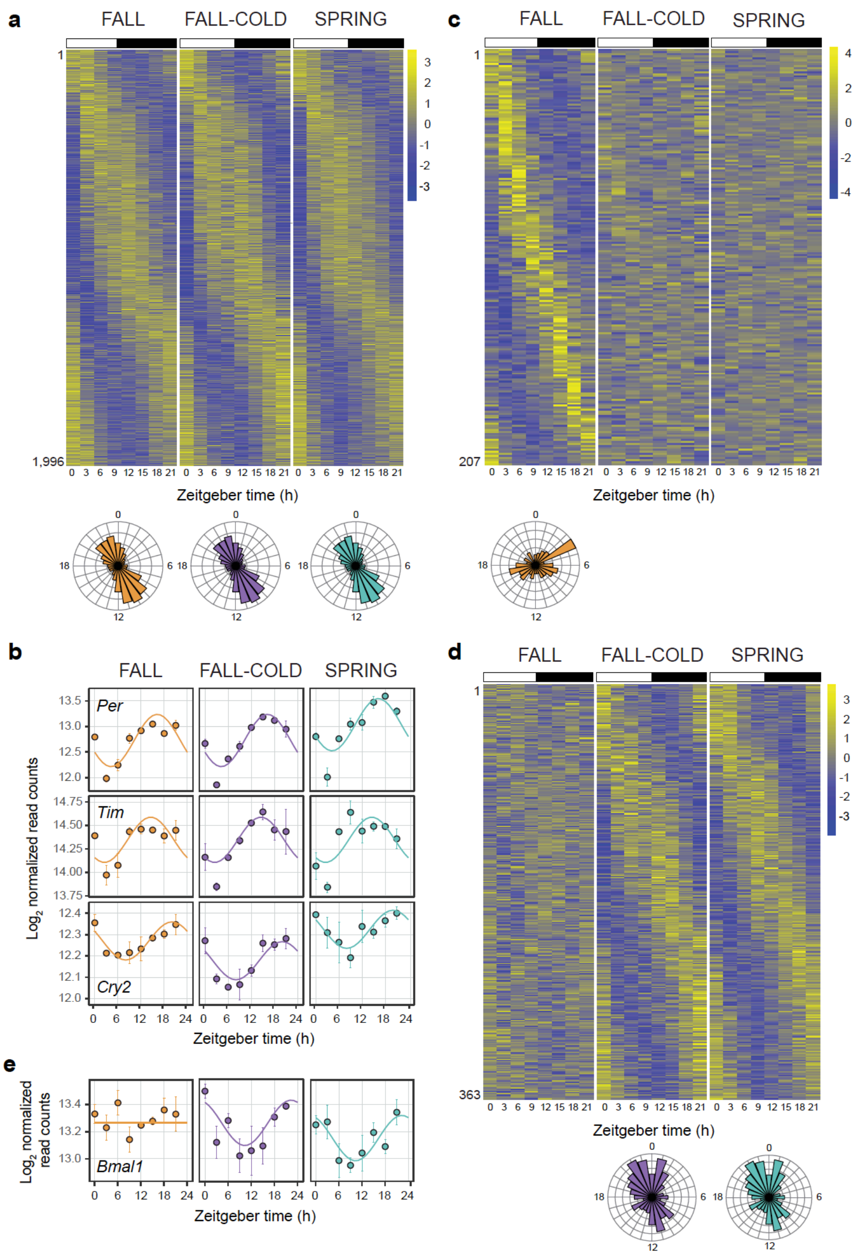


**Supplementary Figure 1: Cold exposure reprograms the rhythmic transcriptome in monarch brains. a**, Heatmaps showing relative RNA levels for genes rhythmically expressed in phase in brains of fall migrants (FALL), cold-treated fall migrants (FALL-COLD) and spring remigrants (SPRING). Columns represent samples collected at different zeitgeber times every 3hr over a 24hr light:dark (LD) cycle. Transcripts are arranged by phase, and their order along the vertical axis is conserved in all conditions. White bars: light; black bars: darkness. See Supplementary Table 4. **b**, $Log_2$ normalized read counts showing temporal expression profiles of the core clock genes *per*, *tim* and *cry2* in brains of all seasonal forms. **c**, Heatmaps showing relative RNA levels for genes rhythmically expressed in brains of fall migrants but arrhythmic in brains of cold-treated fall migrants and spring remigrants. See Supplementary Table 5. **d**,

Heatmaps showing relative RNA levels for genes rhythmically expressed in brains of cold-treated fall migrants and spring remigrants but arrhythmic in brains of fall migrants. See Supplementary Table 6. **e**, $Log_2$ normalized read counts showing temporal expression profiles of the core clock gene *bmal1* in brains of all seasonal forms. For **a**, **c**, and **d**, data represent the mean of 3 replicates of 3 pooled monarch brains per time point for fall and cold-treated fall migrants, and of 4 replicates of 3 pooled monarch brains per time point for spring remigrants. Number of genes is shown at the bottom left of each heatmap. Rose plots at the bottom of the heatmaps depict the phase distribution of rhythmic mRNAs in each condition. For **b** and **e**, each data point represents mean ± s.e.m. of biological replicates and the line shows the fitted curve.

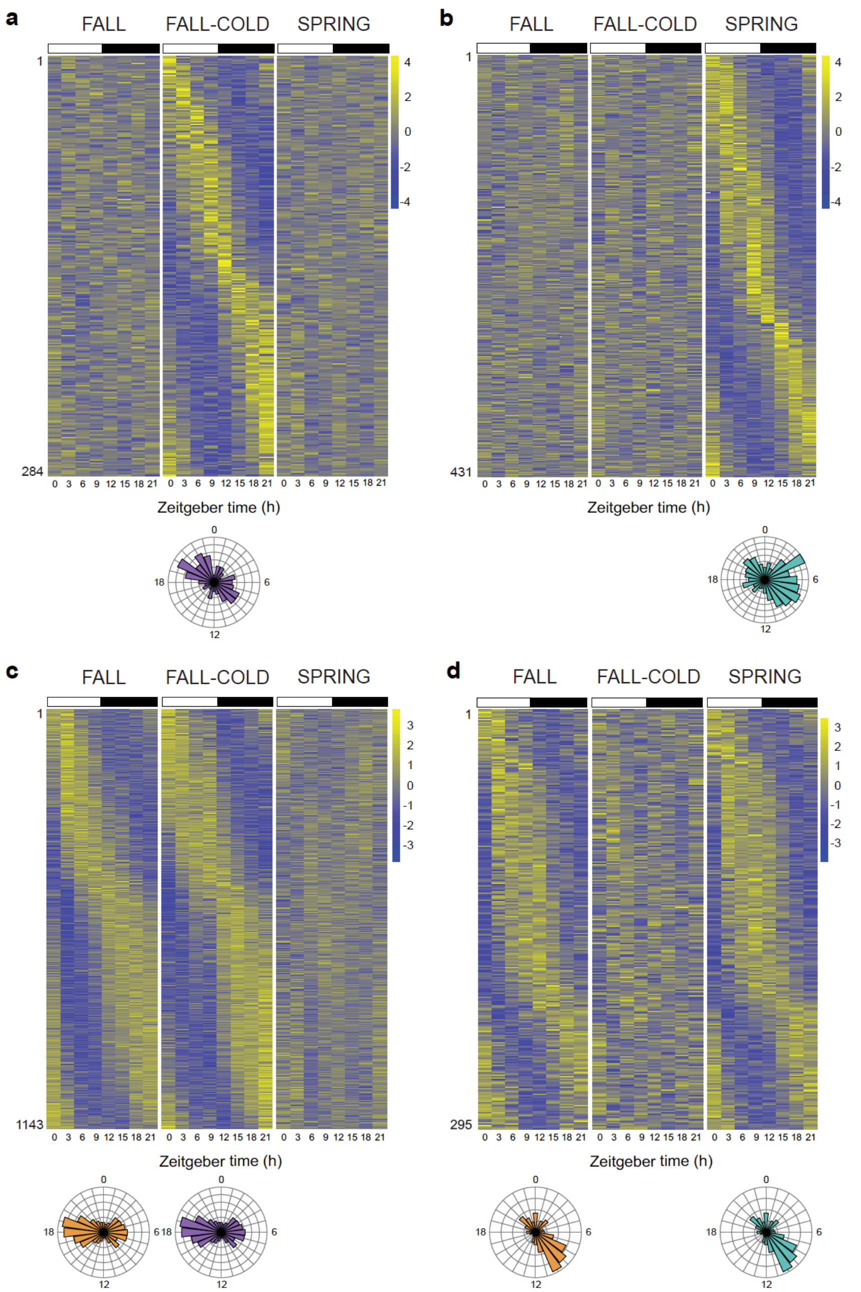


**Supplementary Figure 2: Condition-specific reprogramming of the rhythmic transcriptome in monarch brains. a-d**, Heatmaps showing relative RNA levels for genes rhythmically expressed in brains of a single condition (**a**, **b**) or two conditions (**c**, **d**) not depicted in Figure S1. Columns represent samples collected at different zeitgeber times every 3hr over a 24hr light:dark (LD) cycle. Transcripts are arranged by phase, and their order along the vertical axis is conserved in all conditions. White bars: light; black bars: darkness. Data represent the mean of 3 replicates of 3 pooled monarch brains per time point for fall and cold-treated fall migrants, and of 4 replicates of 3 pooled monarch brains per time point for spring remigrants. Number of genes is shown at the bottom left of each heatmap. Rose plots at the bottom of the heatmaps depict the phase distribution of rhythmic mRNAs in each condition. See Supplementary Tables 7-10.

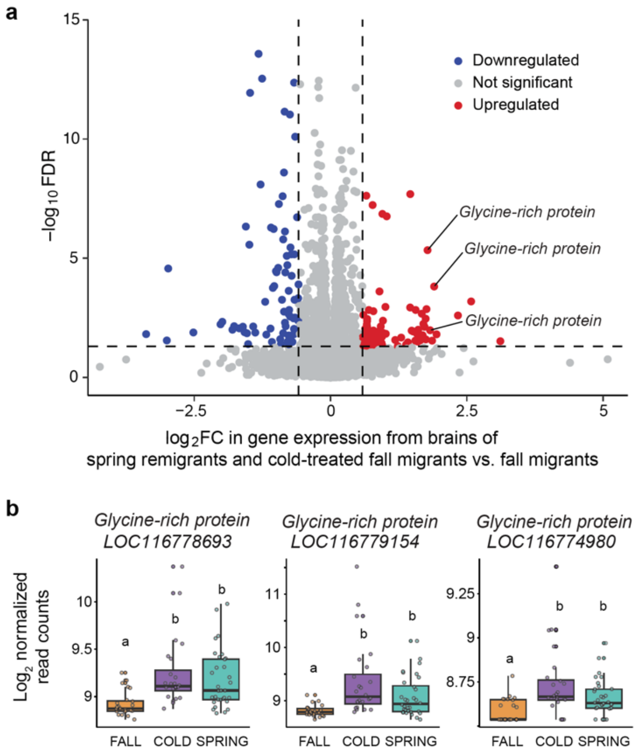


**Supplementary Figure 3: Cold exposure elicits upregulation of *glycine-rich proteins* encoding genes in the monarch brain. a**, Volcano plot depicting differentially expressed genes in brains of spring remigrants/cold-treated fall migrants relative to fall migrants. FC, fold change; FDR, false discovery rate. See Supplementary Table 21. **b**, Box plots and raw data showing the $\log_2$ normalized read counts of expression for 3 *glycine-rich proteins* encoding genes in brains of fall migrants (FALL), cold-treated fall migrants (COLD) and spring remigrants (SPRING) (n = 24 per condition). Error bars on box plots represent 1.5 times the interquartile range. Statistics were obtained by two-tailed Kruskal-Wallis test and post-hoc pairwise comparisons using two-tailed Dunn test with Bonferroni correction. Different letters indicate significant differences between groups ($p < 0.05$). LOC116778693 (Kruskal–Wallis $\chi^2 = 43.28$, $p = 1.60e^{-5}$), LOC116779154 ($\chi^2 = 43.28$, $p = 6.23e^{-6}$), LOC116774980 ($\chi^2 = 43.28$, $p = 4.72e^{-4}$).

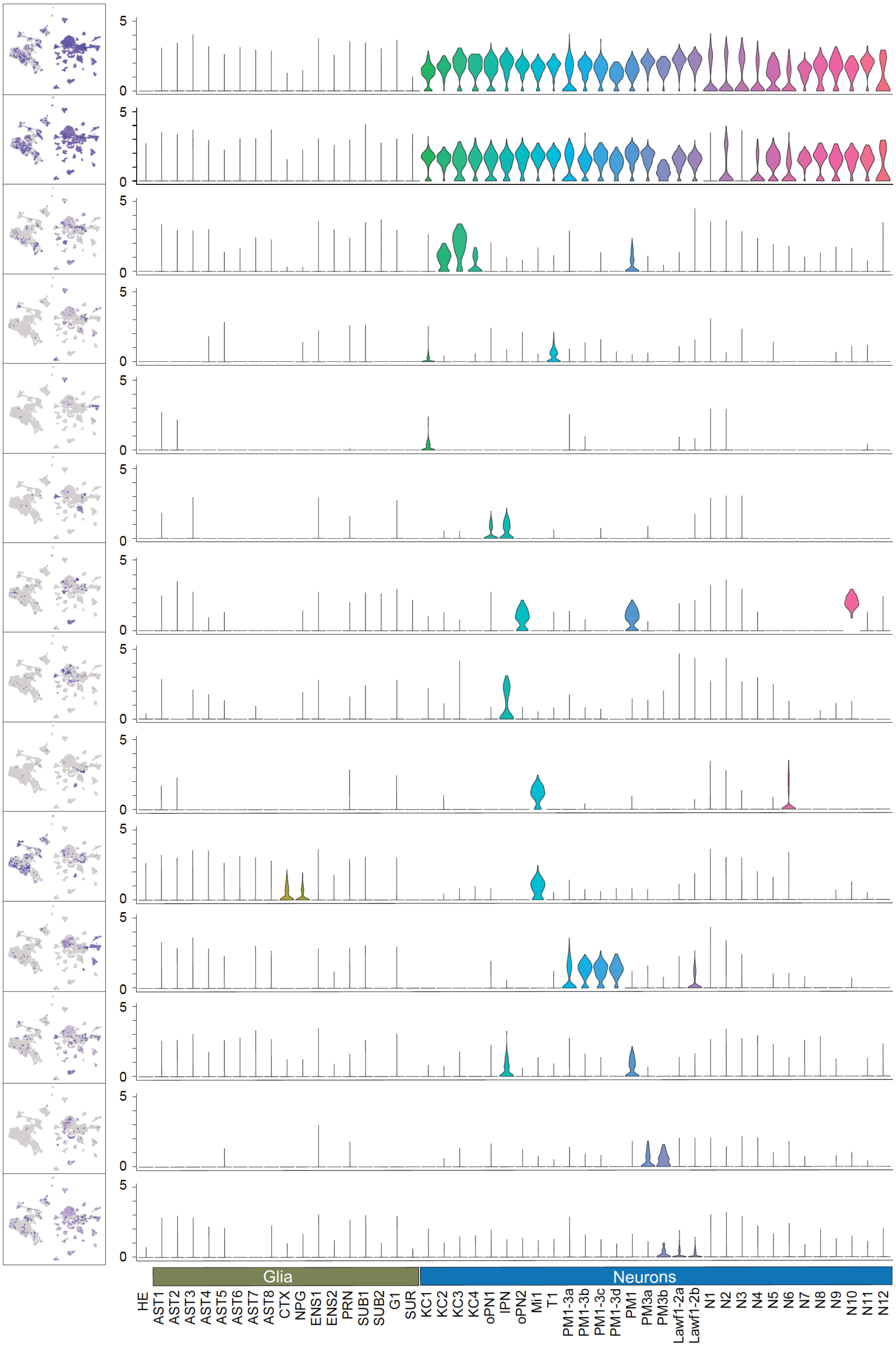
Elav
nSyb
Vmat
Dop1R2
Pax6
Acj6
Vvl
Tk
Bsh
Hth
Lim3
Tiptop
Vsx1
Eya
5
0
Glia
Neurons
HE
AST1
AST2
AST3
AST4
AST5
AST6
AST7
AST8
CTX
NPG
ENS1
ENS2
PRN
SUB1
SUB2
G1
SUR
KC1
KC2
KC3
KC4
oPN1
IPN
oPN2
Mi1
T1
PM1-3a
PM1-3b
PM1-3c
PM1-3d
PM1
PM3a
PM3b
Lawf1-2a
Lawf1-2b
N1
N2
N3
N4
N5
N6
N7
N8
N9
N10
N11
N12

**Supplementary Figure 4: Selected marker genes for annotation of neuronal clusters.** *Left:* Featureplots of marker genes showing per-cell expression levels. *Right*: Violin plots showing expression level per cluster. *Elav*, *Embryonic lethal abnormal visual system*; *nSyb*, *neuronal Synaptobrevin*; *Vmat*, *Vesicular monoamine transporter*; *Dop1R2*, *Dopamine receptor*; *Pax6*, *Paired box 6*; *Acj6*, *Abnormal chemosensory jump 6*; *Vvl*, *Ventral veins lacking*; *Tk*, *Tachykinin*; *Bsh*, *Brain-specific homeobox*; *Hth*, *Homothorax*; *Lim3*, *Lim/homeobox 3*; *Vsx1*, *Visual system homeobox 1*; Eya, *Eyes absent*.

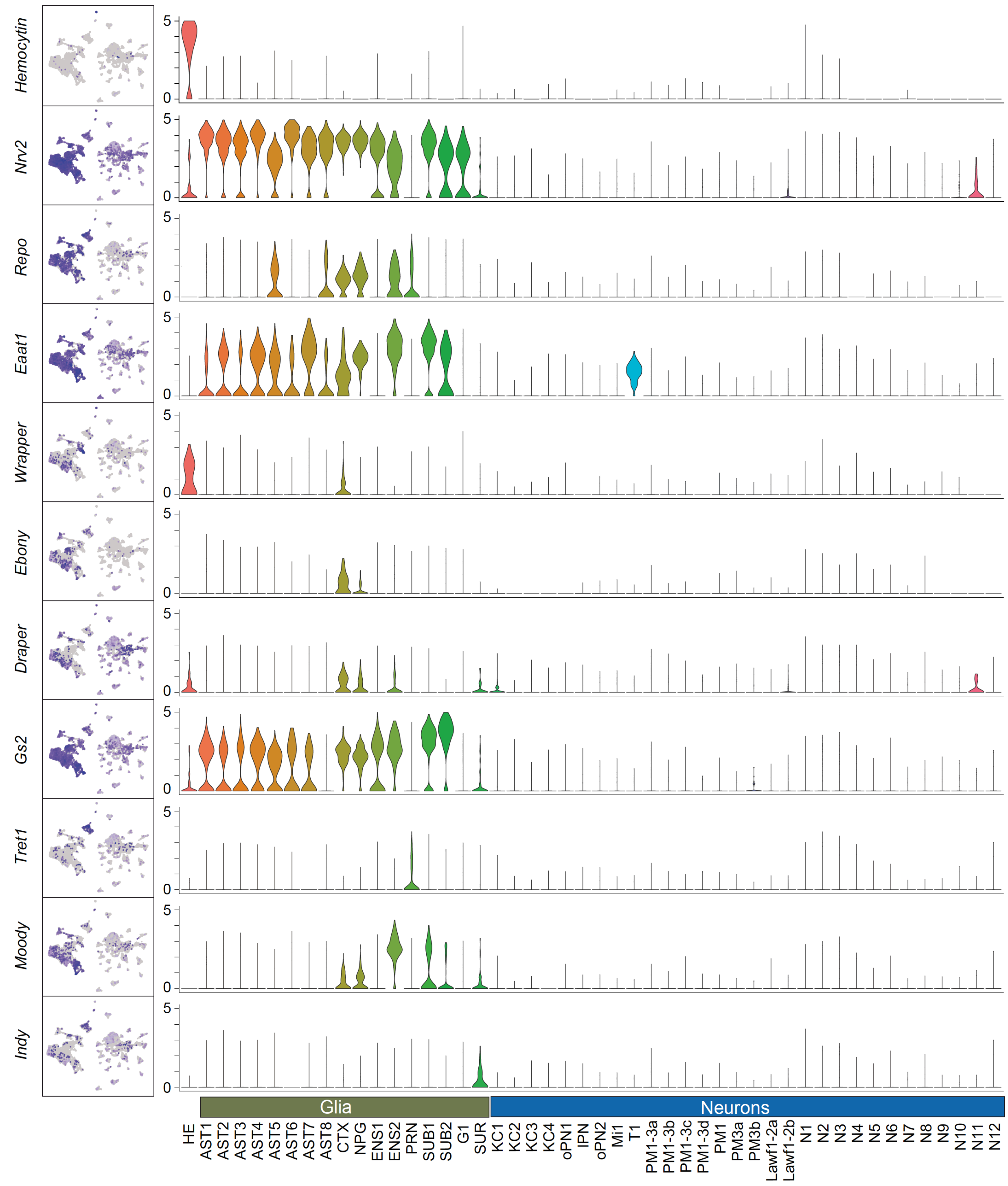


**Supplementary Figure 5: Selected marker genes for annotation of glial clusters.** *Left:* Featureplots of marker genes showing per-cell expression levels. *Right*: Violin plots showing expression level per cluster. *Nrv2*, *Nervana 2*; *Eaat1*, *Excitatory amino acid transporter 1*; *Gs2*, *Glutamine synthetase 2*; *Tret1*, *Trehalose transporter 1*.

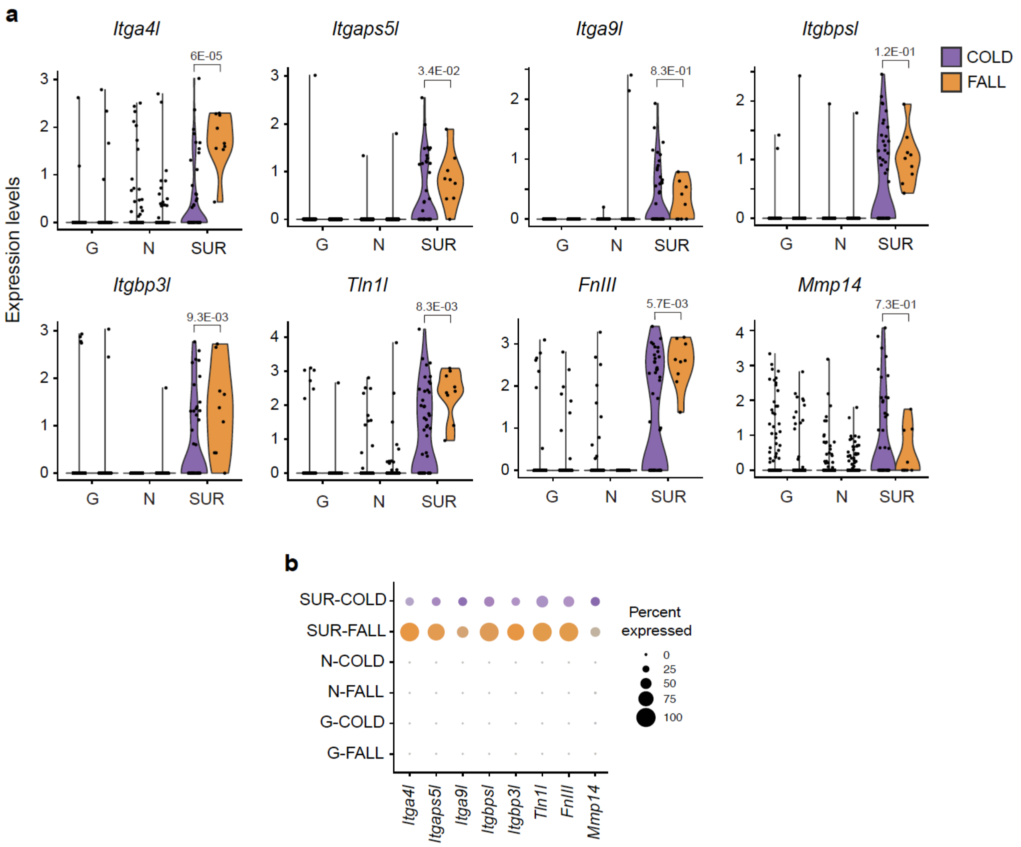


**Supplementary Figure 6: Cold-induced differential expression of genes associated with integrin signaling and ECM composition at single-cell level in surface glia. a**, Violin plots of *Itga4l*, *Itgaps5l*, *Itga9l*, *Itgbpsl*, *Itgbp3l*, *Tln1l*, *FnIII*, *and Mmp14* expression levels in surface glia (SUR), neuronal cell clusters (N), and other glial cell clusters (G) of cold-treated fall migrants and fall migrants. *P*-values from two-tailed Mann Whitney U tests are shown. **b**, Percentage of cells expressing integrins/integrin-related genes in surface glia (SUR) of fall migrants (orange) and cold-treated fall migrants (purple).

**Supplementary Table 1. Summary of RNA-seq pre-processing and mapping for fall migrants (FM)**

| Sample | Total PE reads after quality control | Mapping rate (2nd pass; %) |
|---|---|---|
| FM ZT0 Rep1 | 93,293,998 | 95.32 |
| FM ZT0 Rep2 | 144,559,768 | 95.43 |
| FM ZT0 Rep3 | 125,443,988 | 95.41 |
| FM ZT3 Rep1 | 144,185,130 | 94.76 |
| FM ZT3 Rep2 | 172,796,588 | 97.57 |
| FM ZT3 Rep3 | 130,346,034 | 95.21 |
| FM ZT6 Rep1 | 103,437,360 | 95.24 |
| FM ZT6 Rep2 | 119,681,092 | 96.53 |
| FM ZT6 Rep3 | 110,271,986 | 95.19 |
| FM ZT9 Rep1 | 110,834,428 | 95.35 |
| FM ZT9 Rep2 | 101,932,740 | 95.43 |
| FM ZT9 Rep3 | 107,483,546 | 95.33 |
| FM ZT12 Rep1 | 122,431,570 | 95.14 |
| FM ZT12 Rep2 | 127,002,598 | 94.11 |
| FM ZT12 Rep3 | 117,704,754 | 95.50 |
| FM ZT15 Rep1 | 119,340,716 | 95.08 |
| FM ZT15 Rep2 | 126,364,388 | 95.59 |
| FM ZT15 Rep3 | 53,660,966 | 95.33 |
| FM ZT18 Rep1 | 114,081,286 | 95.05 |

| | | |
|---|---|---|
| FM ZT18 Rep2 | 116,793,086 | 95.47 |
| FM ZT18 Rep3 | 111,340,506 | 95.64 |
| FM ZT21 Rep1 | 86,998,740 | 95.13 |
| FM ZT21 Rep2 | 122,847,702 | 94.12 |
| FM ZT21 Rep3 | 115,984,956 | 95.43 |
| **Average** | **116,617,414** | **95.35** |

**Supplementary Table 2. Summary of RNA-seq pre-processing and mapping for cold-treated fall migrants (CTFM)**

| Sample | Total PE reads after quality control | Mapping rate (2nd pass; %) |
|---|---|---|
| CTFM ZT0 Rep1 | 138,667,532 | 95.40 |
| CTFM ZT0 Rep2 | 116,789,296 | 95.63 |
| CTFM ZT0 Rep3 | 139,414,026 | 95.54 |
| CTFM ZT3 Rep1 | 145,240,640 | 95.16 |
| CTFM ZT3 Rep2 | 127,701,008 | 95.31 |
| CTFM ZT3 Rep3 | 153,252,202 | 95.24 |
| CTFM ZT6 Rep1 | 104,960,634 | 95.20 |
| CTFM ZT6 Rep2 | 108,274,550 | 95.53 |
| CTFM ZT6 Rep3 | 72,765,604 | 94.56 |
| CTFM ZT9 Rep1 | 104,688,390 | 95.68 |
| CTFM ZT9 Rep2 | 100,689,506 | 95.24 |
| CTFM ZT9 Rep3 | 127,985,998 | 95.14 |
| CTFM ZT12 Rep1 | 126,783,532 | 95.48 |
| CTFM ZT12 Rep2 | 104,275,468 | 95.37 |
| CTFM ZT12 Rep3 | 129,535,040 | 90.04 |
| CTFM ZT15 Rep1 | 115,151,644 | 95.47 |
| CTFM ZT15 Rep2 | 116,934,944 | 95.37 |
| CTFM ZT15 Rep3 | 137,830,706 | 95.48 |

| | | |
|---|---|---|
| CTFM ZT18 Rep1 | 122,645,154 | 93.47 |
| CTFM ZT18 Rep2 | 113,854,730 | 95.46 |
| CTFM ZT18 Rep3 | 135,516,766 | 95.70 |
| CTFM ZT21 Rep1 | 92,998,002 | 93.41 |
| CTFM ZT21 Rep2 | 84,325,722 | 95.51 |
| CTFM ZT21 Rep3 | 139,714,484 | 88.36 |
| **Average** | **119,166,482** | **94.70** |

**Supplementary Table 3. Summary of RNA-seq pre-processing and mapping for spring remigrants (SR)**

| Sample | Total PE reads after quality control | Mapping rate (2nd pass; %) |
|---|---|---|
| SR ZT0 Rep1 | 160,711,306 | 89.39 |
| SR ZT0 Rep2 | 180,403,350 | 94.08 |
| SR ZT0 Rep3 | 216,241,976 | 93.53 |
| SR ZT0 Rep4 | 174,531,384 | 93.66 |
| SR ZT3 Rep1 | 196,381,830 | 93.67 |
| SR ZT3 Rep2 | 201,494,910 | 95.22 |
| SR ZT3 Rep3 | 228,844,048 | 93.04 |
| SR ZT3 Rep4 | 198,910,580 | 92.98 |
| SR ZT6 Rep1 | 160,285,470 | 94.83 |
| SR ZT6 Rep2 | 192,311,310 | 95.44 |
| SR ZT6 Rep3 | 204,832,734 | 94.09 |
| SR ZT6 Rep4 | 177,593,560 | 95.06 |
| SR ZT9 Rep1 | 157,160,550 | 95.06 |
| SR ZT9 Rep2 | 182,386,078 | 95.39 |
| SR ZT9 Rep3 | 191,029,928 | 95.01 |
| SR ZT9 Rep4 | 173,669,086 | 94.76 |
| SR ZT12 Rep1 | 181,657,474 | 94.79 |
| SR ZT12 Rep2 | 190,230,718 | 94.75 |

| | | |
|---|---|---|
| SR ZT12 Rep3 | 213,955,824 | 95.03 |
| SR ZT12 Rep4 | 183,095,288 | 94.33 |
| SR ZT15 Rep1 | 192,548,752 | 93.69 |
| SR ZT15 Rep2 | 189,684,324 | 95.10 |
| SR ZT15 Rep3 | 215,439,270 | 94.69 |
| SR ZT15 Rep4 | 180,672,180 | 94.48 |
| SR ZT18 Rep1 | 171,912,744 | 93.22 |
| SR ZT18 Rep2 | 187,892,496 | 94.97 |
| SR ZT18 Rep3 | 181,206,458 | 95.12 |
| SR ZT18 Rep4 | 178,280,270 | 93.57 |
| SR ZT21 Rep1 | 159,567,622 | 93.06 |
| SR ZT21 Rep2 | 193,705,104 | 95.06 |
| SR ZT21 Rep3 | 186,336,460 | 95.24 |
| SR ZT21 Rep4 | 174,660,842 | 94.12 |
| **Average** | **186,801,060** | **94.26** |

**Supplementary Tables 4 to 22 are in excel files format.**
